# *STEMorph*: A Set of Morphed Emotional Face Stimuli from Angry to Happy Derived from *NimStim*

**DOI:** 10.1101/2024.05.13.593881

**Authors:** Mohammad Ebrahim Katebi, Mohammad Hossein Ghafari, Tara Ghafari

## Abstract

Emotion recognition through facial expressions is crucial for interpreting social cues. However, it is often influenced by biases, i.e., systematic recognition advantages for particular emotions. Nevertheless, these biases are inconsistently reported across studies, likely due to methodological variations which underline the necessity for a standardized approach. Traditional face morphing methods can create unnatural-looking stimuli, and may confound the interpretation of emotions. To address this issue, we here introduce *STEMorph*, a validated stimulus set based on the *NimStim* set. We employed neutral-anchored morphing and neural-network-generated masks to reduce morphing artifacts and preserve the coherence of the depicted expressions. We validated our stimulus set by having participants rate each face on a 9-point scale ranging from angry to happy, assessing the perceived emotional intensity. *STEMorph’s* validity was confirmed through linear mixed-effects modelling, showing a strong association between subjective ratings and intended morph level while accounting for other effects. Moreover, aligning with previous research highlighting gender as a key factor in emotion recognition, *STEMorph* also showed variation across participant-gender dimension. *STEMorph’s* reliability was confirmed through a two-week follow-up rating session with a subgroup of the same participants. By introducing a controlled and empirically evaluated stimulus set of morphed emotional faces, *STEMorph* provides a useful resource for future investigations of facial emotion recognition.

## Introduction

Emotion recognition is a fundamental aspect of human social interaction, allowing individuals to interpret and respond appropriately to the emotional states of others. Facial expressions serve as a primary modality for conveying emotions, providing valuable cues about an individual’s internal states (1). However, despite its significance, the process of emotion recognition is not devoid of biases. These biases manifest as systematic recognition advantages for particular emotions and have significant implications for both clinical and non-clinical populations (2).

Recognition biases can vary across different conditions. Individuals with major depressive disorder (MDD) tend to interpret neutral or ambiguous faces as sad or threatening, reflecting a negativity bias. This aligns with findings that people with MDD are more attuned to negative emotions and less responsive to positive ones, which impairs their ability to interpret emotional cues (3, 4). Such biases may reinforce mood-congruent cognitive distortions, contributing to the maintenance and exacerbation of depressive symptoms (5, 6). Similarly, individuals with generalized anxiety disorder (GAD) and social anxiety disorder (SAD) exhibit hypervigilance toward negative emotional cues, such as anger or fear, which can lead to misinterpretations of facial expressions and reinforce maladaptive emotional responses (7, 8). Therefore, these biases can significantly impact social interactions and clinical outcomes; emphasizing the importance of their precise measurement for both research and clinical applications.

Studies often report inconsistent results regarding the direction and magnitude of emotional recognition biases (9). For instance, one study found that healthy individuals did not demonstrate a bias toward happiness or sadness when presented with neutral facial expressions (10). While another study showed that healthy participants tended to perceive neutral faces as expressing happiness, whereas depressed individuals interpreted them as neutral (8). It is uncertain whether these mixed findings reflect genuine individual differences in emotion perception or are due to methodological variations between the studies (11). Particularly, the quality and ecological validity of facial stimuli can significantly impact participants’ responses. While standardized photographic stimuli are commonly used for their control over visual properties, they may not fully capture the complexities of real-world facial expressions, which can lead to discrepancies in findings. Research indicates that more naturalistic stimuli, such as dynamic or spontaneous facial expressions, enhance ecological validity but introduce variability that complicates experimental control (12, 13). Consequently, a key challenge lies in creating stimuli that maintain natural appearance while allowing precise control over emotional expressions (12, 13).

One widely used approach to generating a continuum of emotions is through face morphing techniques. Face morphing is an image processing technique employed to achieve transformation from one face to another and facilitates a gradual transition between two facial images. By morphing facial features between two extreme emotional expressions, researchers generate a spectrum of face images with ambiguous emotional expressions for participants to interpret (14, 15). However, traditional face morphing presents challenges, particularly concerning the formation of unrealistic and unnatural facial stimuli (15) which may confound interpretations and obscure real emotion recognition processes.

To reduce artifacts introduced by morphing in non-face regions, careful masking of the face images (extracting the face area while removing the background) is required. Using no mask (full head) or simple oval/square masks which are employed by many studies (12) may cause distortions that risk emotional content loss, since various facial features hold different degrees of significance depending on the type of expression (16, 17). For instance, the mouth region is particularly salient for recognizing happiness, while the eyes are more critical for identifying fear or sadness (18). These regional differences can lead to biases in emotion perception, as individuals may focus on different facial features based on cultural or individual preferences (17). These underscore the necessity for stimuli preserving anatomical fidelity through methods like neutral-anchored morphing (11) and standardized masking. However, there are no existing publicly available stimulus set without these methodological confounds (12, 15, 19). While sets like the Karolinska Directed Emotional Faces (KDEF) (20) and the Pictures of Facial Affect (POFA) (21) provide valuable resources, they often rely on basic morphing techniques that can produce artifacts, particularly around facial features crucial for emotion recognition such as the eyes and mouth. Furthermore, many existing sets use simple geometric masks (oval or rectangular), which are practical but may introduce artificial mask boundaries or remove facial regions unevenly across identities (12, 15). To address this gap, we developed and validated a new stimulus set utilizing neutral-anchored morphing alongside neural-network-generated masks, to optimize naturalism and preserving perceptual coherence across the continuum.

We derived the stimuli from the *NimStim set of facial expressions* (22) which is a comprehensive collection of facial images portraying a spectrum of emotions including happy, angry, and neutral expressions. The selection of anger and happiness as the target emotions for this study was based on their distinct valence and arousal profiles, which make them suitable for exploring biases in emotion recognition. Additionally, these emotions are commonly used in emotion research (2, 11), facilitating comparisons with existing literature. Investigating the transition between these emotions allows us to examine biases in emotional perception across a clear positive-negative emotional gradient, which is particularly relevant for studies on mood disorders and social cognition (2). We adopt a nuanced approach to morphing emotional faces by generating distinct naturalistic face masks for each image using neural networks. Furthermore, we use the neutral facial expression as a central anchor point to improve the structural coherence of the transition between anger and happiness. We then evaluate the validity and reliability of *STEMorph* by asking participants to rate the emotional content of these stimuli. In doing so, the present paper has a dual but unified purpose: to introduce *STEMorph* as a publicly available stimulus set designed for research on facial emotion recognition, and to report the psychophysical validation that establishes its perceptual coherence.

Importantly, prior literature indicates that both the gender of the face and the gender of the perceiver influence emotion recognition. Female faces have been consistently rated as more emotionally expressive and displaying greater happiness than comparable male faces (17, 23). Simultaneously, female perceivers tend to demonstrate greater sensitivity to emotional gradations in facial stimuli, yielding steeper psychometric slopes when rating morphed expressions (24). Including both face gender and participant gender in our validation framework therefore serves two purposes: it enables convergent validity testing by checking whether *STEMorph* reproduces established gender-related effects, and it provides normative reference data to facilitate comparisons across studies using different participant demographics.

## Method

### Participants

A total of 50 participants (age range: 18–26 years; mean age: 21.5; 22 females) were recruited for the validity testing. All participants were medical students from either Tehran University of Medical Sciences (TUMS) or Shahid Beheshti University of Medical Sciences (SBUMS). All participants reported having normal or corrected-to-normal vision and no history of diagnosed neurological or psychiatric disorders. Written informed consent was obtained from each participant prior to participation, and all were compensated financially for their time. The study received ethical approval from the Ethics Committee of Shahid Beheshti University of Medical Sciences.

### Experiment design & Face Generation

We used face morphing to produce a linear transition of emotions from anger to happiness using neutral expression as an intermediary point. The stimuli were derived from the *NimStim set of facial expressions* (*22*). Each emotional expression has an open-mouth and closed-mouth variation in *NimStim*, from which we selected closed-mouth variations. Closed-mouth variants were selected to reduce variability introduced by visible teeth, tongue, and large mouth-aperture changes, which can influence perceived happiness and may introduce additional identity-specific artifacts during morphing. This choice improved experimental control but also restricts the generalisability of the stimulus set to closed-mouth expressions. *Abrosoft Fantamorph* (*25*) software was used for morphing, involving nine steps to transition from the angry face to neutrality and subsequently to the happy face. We used the original facial expressions of anger, happiness, and neutral as reference points and generated three intermediate emotional states between each original face.

We selected 22 *NimStim* identities (11 female and 11 male) based on predefined technical and perceptual criteria. Included identities had to provide usable closed-mouth angry, neutral, and happy expressions; comparable head pose, gaze direction, framing, and illumination across the three anchor images; no major occlusion of internal facial features; and sufficient landmark alignment to allow a visually continuous angry-neutral-happy morph sequence. Identities were excluded when the expression intensity or facial configuration produced disruptive feature misalignment, visible warping, or mask-boundary artifacts. The final set is composed of 198 distinct faces, representing 9 different emotional steps for each of the 22 individuals (Figure 1).

**Figure 1.**
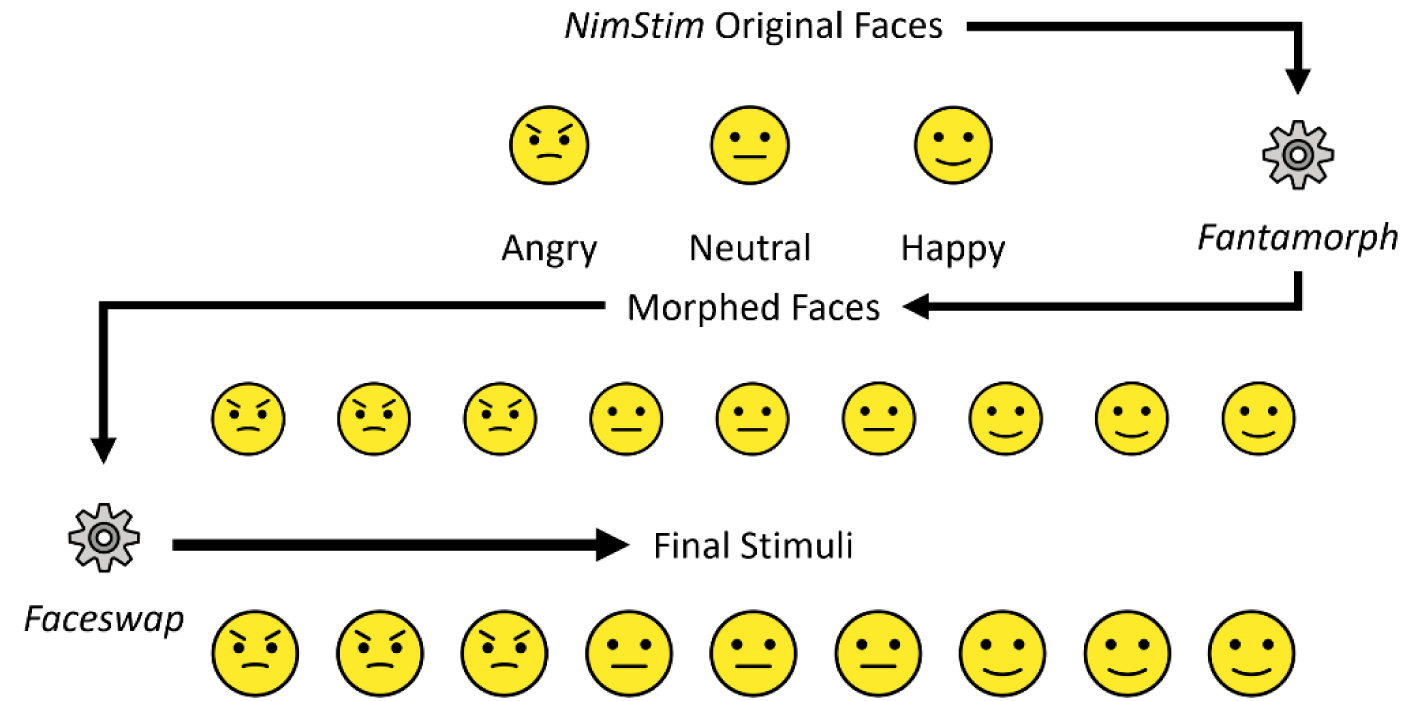
Example Stimuli from the Set. Note. Top row illustrates the two ends of the spectrum (happy and angry) along with the middle step (neutral). Middle row shows the intermediary morphing stages without masking. Bottom row shows the final outcome with corresponding neural-network-generated masks applied to each morphing step.

To address artifacts introduced in the ears, hair, and other non-landmark regions of the face during conventional face morphing, we generated distinct naturalistic face masks for each image using the *FaceSwap* (*26*) software, which utilizes the *BiSeNet-FP* (*27*) neural network. *BiSeNet-FP* is a modified lightweight neural network designed for face parsing. The model was set to not include hair or ears and extract only the face region. Manual editing was restricted to correcting mask-boundary errors, such as splits in the face region caused by stray hair, or erroneous inclusion of hair, ears, or background. Corrections were standardized by applying the same rule to all stimuli: the final mask had to include the continuous visible face region while excluding hair, ears, and background.

### Experimental Setup

Participants were asked to rate the emotional state of each presented stimulus on a 9-point scale from angry to happy. The task was developed and implemented using *Psychtoolbox* (28) within *MATLAB R2022a* (Figure 2). Participants were comfortably seated in a dimly lit room facing a laptop monitor. The resolution of monitor screen was 1920 × 1080 and it was positioned ∼54 cm from the participant. Participants were instructed to freely observe the images without fixation requirements.

**Figure 2.**
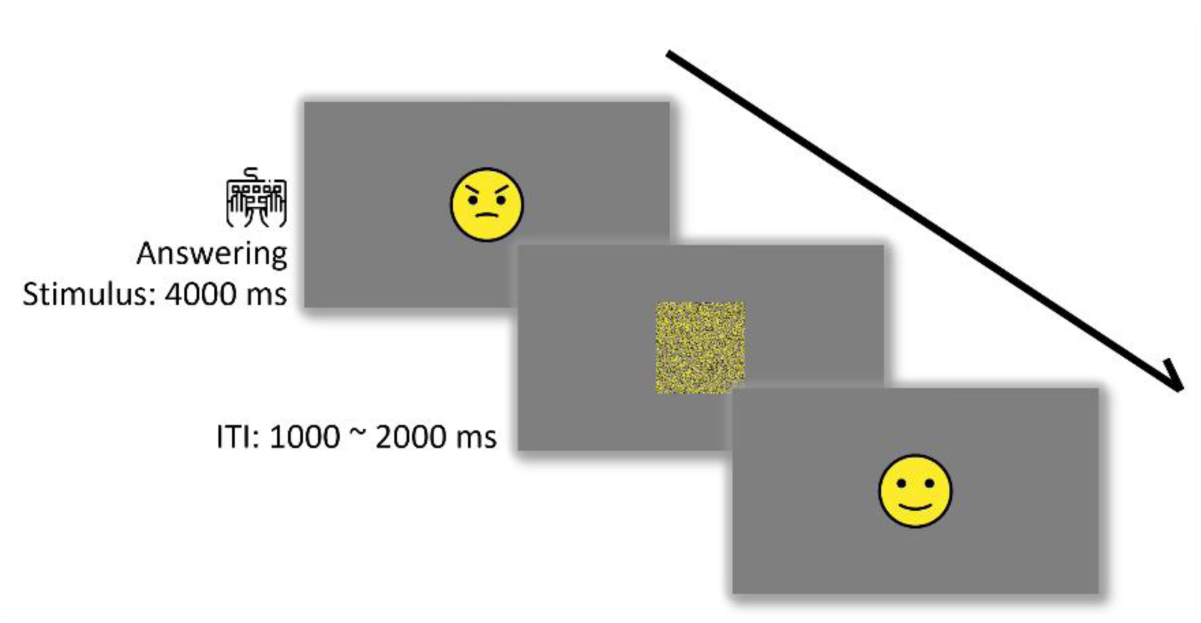
Schematic presentation of the experimental design. Note. Stimuli appeared at the center of the screen, and participants rated the emotional intensity of each face within a 4000 ms viewing window without fixation restrictions. Each stimulus was followed by a randomized inter-trial interval (ITI) of 1000 to 2000 ms, during which a noise mask was displayed.

**Figure 3.**
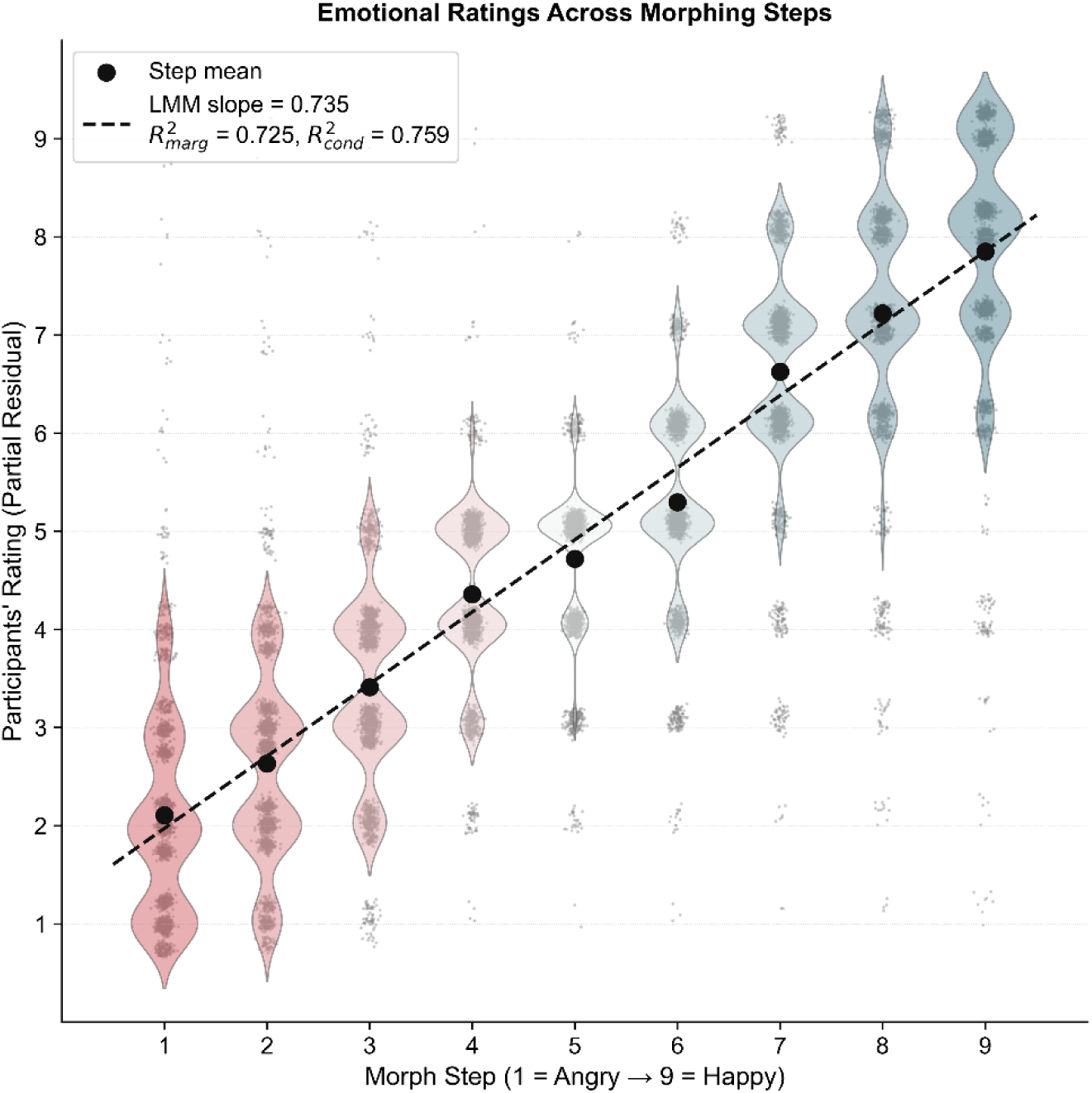
Validity of emotional ratings across morphing steps. Note. Morphing steps range from angry (1) to happy (9) on the x-axis. The y-axis shows participants’ emotional ratings, adjusted for face gender, participant gender, their interactions, and random effects of participant and face identity using a mixed-effects model. Violin plots illustrate the distribution of ratings at each morph step. For visual clarity, extreme outliers (±2 SD from the mean rating at each morph step) were excluded from the plot, although all data were included in the statistical analyses. Small grey dots represent individual trial ratings after adjustment for model covariates, large black dots indicate mean ratings, and the dashed line shows the fitted mixed-effects model trend. The figure also reports marginal R² (variance explained by fixed effects) and conditional R² (variance explained by both fixed and random effects).

The task started with pressing a key, after which participants were asked to rate the emotional state of each presented face based on a 9-point scale where 1 indicated the angriest emotional state and 9 indicated the happiest emotional state. The exact instructions were: ‘Please rate the emotional state of each face on a scale from 1 to 9, where 1 represents the angriest expression and 9 represents the happiest expression.’ To control for motor and numerical biases, the scale was reversed for half of the participants. For these individuals, the instructions were adjusted accordingly, and their responses were rescaled during analysis to align with the original scale (1 = angry, 9 = happy). Each participant had a 4-second window to provide their rating; in the absence of a response within this timeframe, the task automatically proceeded to the subsequent face.

Each of the 198 stimuli was presented twice in a randomized order, resulting in a total of 396 trials per participant, interspersed by two 2-minute breaks. Following each trial, a randomized inter-trial interval (ITI) of 1000 to 2000 ms ensued, during which a noise mask was displayed at the center of the screen to prevent from interference between trials and minimize priming effects (29, 30). To generate the noise mask the stimulus presented in the previous trial was scrambled into 4-pixel squares.

### Reliability Assessment

To assess reliability, the same behavioral task was readministered to a subgroup of 23 participants two weeks following their initial participation. This subgroup consisted of participants who agreed to return for retesting (age range: 18-26, mean age: 22, 5 females).

### Data Analysis

#### Validity

We used a linear mixed-effects model to investigate the relationship between morph level and subjective ratings implemented using *Statsmodels* (31). The model included fixed effects of morph step (1-9), face gender (0 = female, 1 = male), participant gender (0 = female, 1 = male), and all two-way interactions. To account for clustering of repeated observations, random intercepts were included for participant and face identity. The model was specified as:

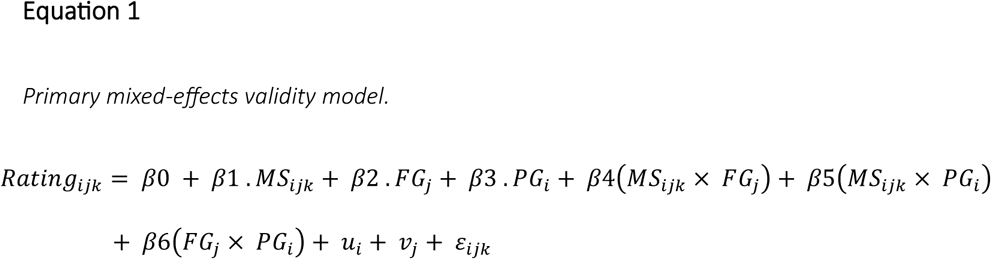

where *i* indexes participant, *j* indexes face identity, and *k* indexes trial/observation. *MS* corresponds to Morph Step, *FG* to Face Gender, and *PG* to Participant Gender. The terms *μ*_*i*_ and *ν*_*j*_ represent random intercepts for participant and face identity, respectively, and *ε*_*ijk*_ represents the residual error. To visualize partial regression plots we generated violin plots using *Matplotlib* (32) and *Seaborn* (33).

### Gender Subgroup Analysis

To examine gender-specific patterns in emotion perception, we derived participant-gender and face-gender subgroup visualisations from the full model. This approach preserved the full structure while allowing the morph-rating relationship to be visualised within each gender dimension. For participant-gender subgroup visualisation, we computed partial residuals by removing the fixed-effect contribution of face gender terms from the full model:

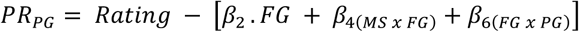

For face-gender subgroup visualization, we computed partial residuals by removing the fixed-effect contribution of participant gender terms from the full model:

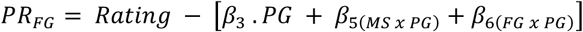

### Reliability

To evaluate test-retest reliability, we analysed data from participants who completed the retest session two weeks later. Trials were matched across sessions based on participant, face identity, morph step, and repetition. The primary reliability metric was Intraclass Correlation Coefficients (ICC) (2,1), a two-way random-effects, absolute-agreement, single-measure intraclass correlation coefficient calculated using *Pingouin* (34). This metric was selected because it evaluates whether ratings agree in absolute value across the two sessions.

## Results

The aim of this study was to develop and validate a new stimulus set for emotion recognition utilizing neutral-anchored face morphing and neural-network-generated masks to optimize naturalism and coherence. A total of 50 participants were recruited to rate the emotional content of the face stimuli on a 9-point scale ranging from angry to happy with a subset of 23 participants returning for a reliability assessment two weeks later.

### Validity

We used a linear mixed-effects model to examine emotional ratings (mean = 4.85, SD = 2.14) as a function of morph step while accounting for face gender, participant gender, their two-way interactions, and random intercepts for participant and face identity (Table 1). Morph step strongly predicted emotional ratings (β= 0.735, SE = 0.005, 95% CI [0.725, 0.745], *p* < 0.001), indicating that participants’ ratings closely tracked the intended anger-to-happiness continuum. The fixed effects explained a large proportion of variance (marginal R^2^ = 0.7251), and the full model including random effects explained slightly more variance (conditional R^2^ = 0.7591).

**Table 1.**
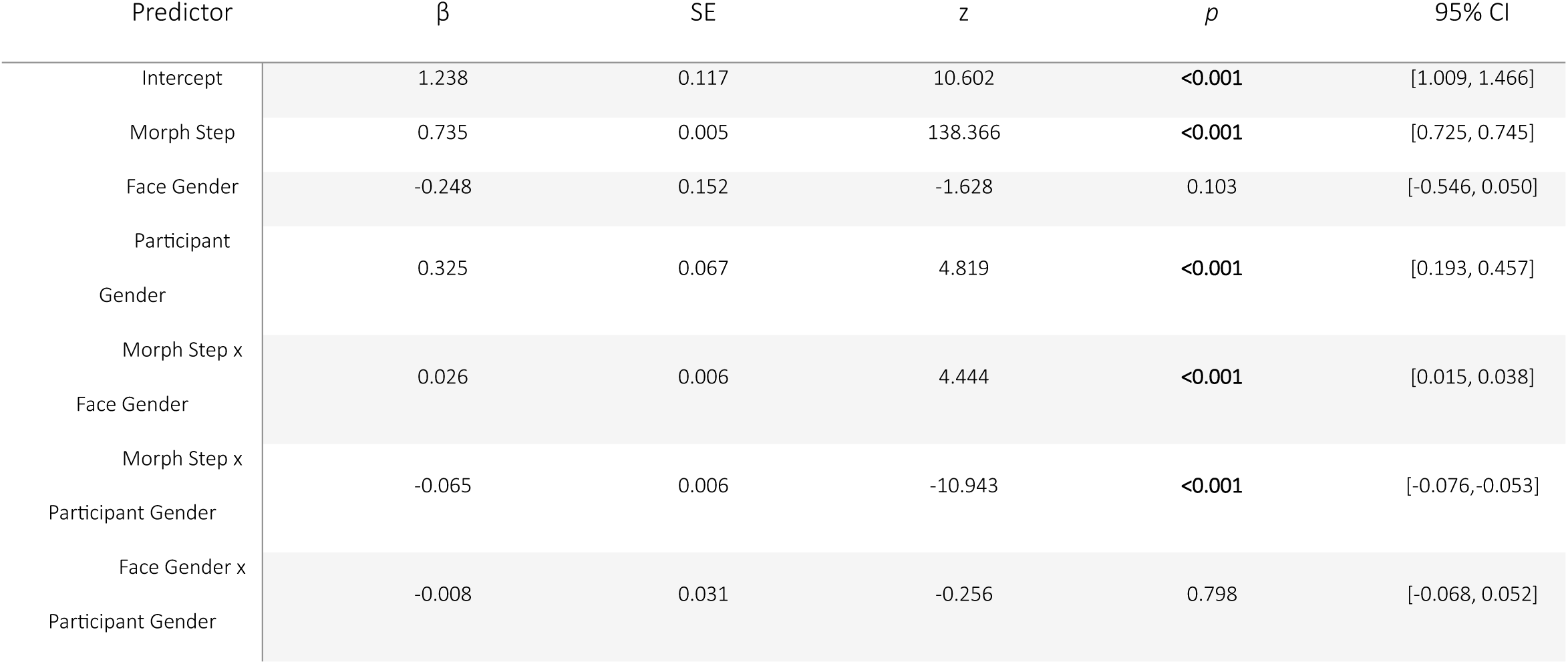
Linear Mixed-Effects Model Predicting Emotional Ratings. Note. Linear mixed-effects model predicting emotional ratings. The model included fixed effects of morph step, face gender, participant gender, and all two-way interactions. Morph step represents emotion intensity progression from anger (1) to happiness (9). Face gender and participant gender were coded 0 = female and 1 = male.

| Predictor | $\beta$ | SE | z | p | 95% CI |
| --- | --- | --- | --- | --- | --- |
| Intercept | 1.238 | 0.117 | 10.602 | <0.001 | [1.009, 1.466] |
| Morph Step | 0.735 | 0.005 | 138.366 | <0.001 | [0.725, 0.745] |
| Face Gender | -0.248 | 0.152 | -1.628 | 0.103 | [-0.546, 0.050] |
| Participant Gender | 0.325 | 0.067 | 4.819 | <0.001 | [0.193, 0.457] |
| Morph Step x Face Gender | 0.026 | 0.006 | 4.444 | <0.001 | [0.015, 0.038] |
| Morph Step x Participant Gender | -0.065 | 0.006 | -10.943 | <0.001 | [-0.076, -0.053] |
| Face Gender x Participant Gender | -0.008 | 0.031 | -0.256 | 0.798 | [-0.068, 0.052] |

Across the full design, 19,800 trials were expected. A total of 204 trials timed out without a response, corresponding to 1.03% of all trials and leaving 19,596 valid ratings for the primary mixed-effects model. Timed-out trials were not assigned ratings and were excluded from all analyses. The rate of these responses varied slightly across morph steps (χ² = 15.69, p = 0.047) and was associated with participant gender (χ² = 7.46, p = 0.006), but did not differ significantly by face gender (χ² = 3.10, p = 0.079). Because the overall loss is only about 1% and timed-out trials carry no rating, this is unlikely to have biased the central findings.

In a validation study, extreme but genuine ratings are themselves part of the perceptual response we are trying to characterise and discarding them risks presenting the set as more orderly than it is; we therefore retained the full dataset for the primary mixed-effects model. To demonstrate that our conclusions do not depend on this choice, we conducted a sensitivity analysis in which ratings more than two standard deviations from the morph-step mean were removed (1,048 observations, 5.35%) and the model was refit. The morph-step slope increased only slightly (from β = 0.735 to β = 0.771, p < 0.001), and the direction and significance of every effect were preserved.

Participants generally responded well before the 4-second deadline. Across valid trials, mean response time was 1.827s (median = 1.695s, SD = 0.695s). In a mixed-effects model of log-response time, there was no significant linear effect of morph step (p = 0.174), but a small, statistically significant quadratic (inverted-U pattern) effect (p = 0.003): response times were marginally longer for the more ambiguous central morph steps than for the emotional extremes. Given that responses were typically made within approximately two seconds, well below the maximum response window, the 4-second limit functioned primarily as a response deadline, and the slight central slowing is consistent with greater perceptual ambiguity near the neutral anchor (35).

### Gender Subgroup Validity Analysis

To assess the robustness of validity across gender dimensions, we included face gender and participant gender in the mixed-effects model described above (Equation 1). Significant effects were observed for participant gender (β= 0.325, *p* < 0.001), the interaction between morph step and face gender (β= 0.026, *p* < 0.001), and the interaction between morph step and participant gender (β=-0.065, *p* < 0.001; see Figure 4). The main effect of face gender was not significant after accounting for face identity (β=-0.248, *p* = 0.103), and the face gender x participant gender interaction was also not significant (β=-0.008, *p* = 0.798).

**Figure 4.**
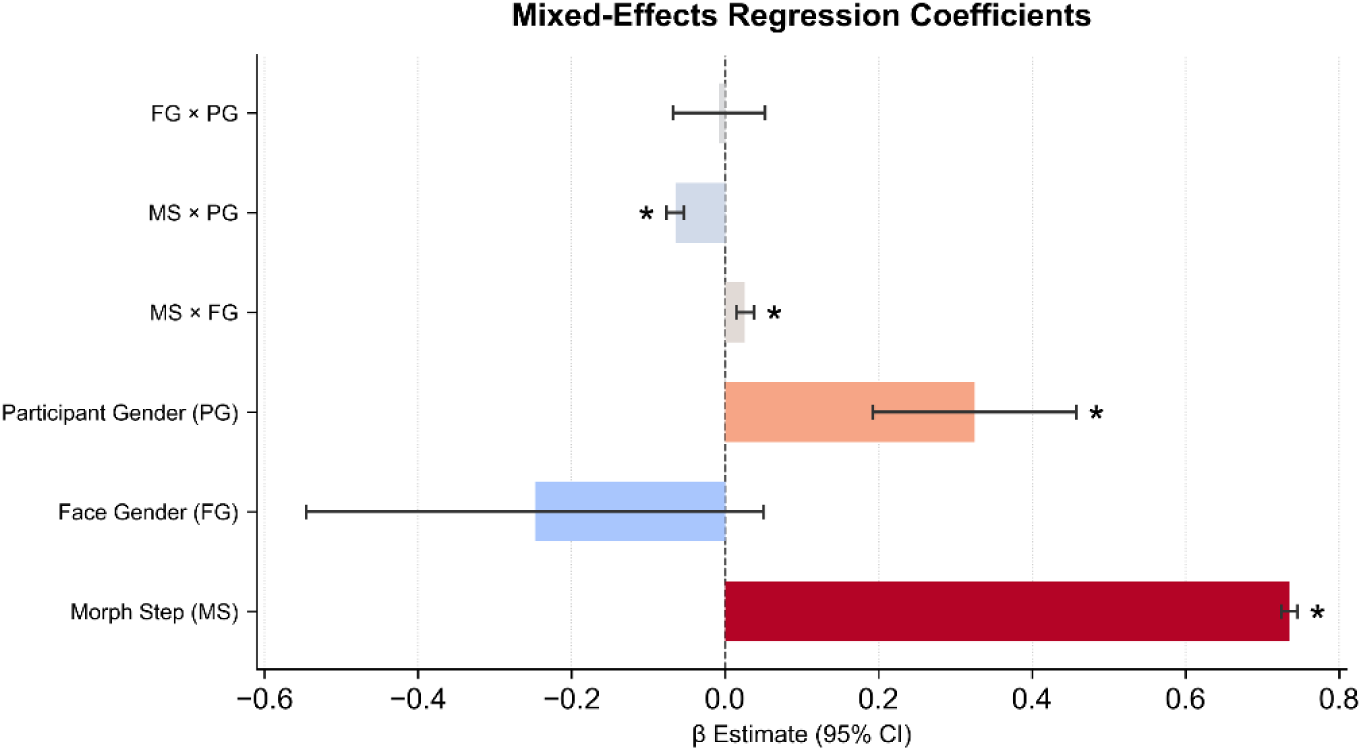
Regression coefficients for the model. Note. Coefficient estimates and 95% CIs from the linear mixed-effects model. Predictors include Morph Step (MS; 1–9), Face Gender (FG; 0 = female, 1 = male), Participant Gender (PG; 0 = female, 1 = male), and their two-way interactions. Asterisks denote significancy at α = 0.05.

**Figure 5.**
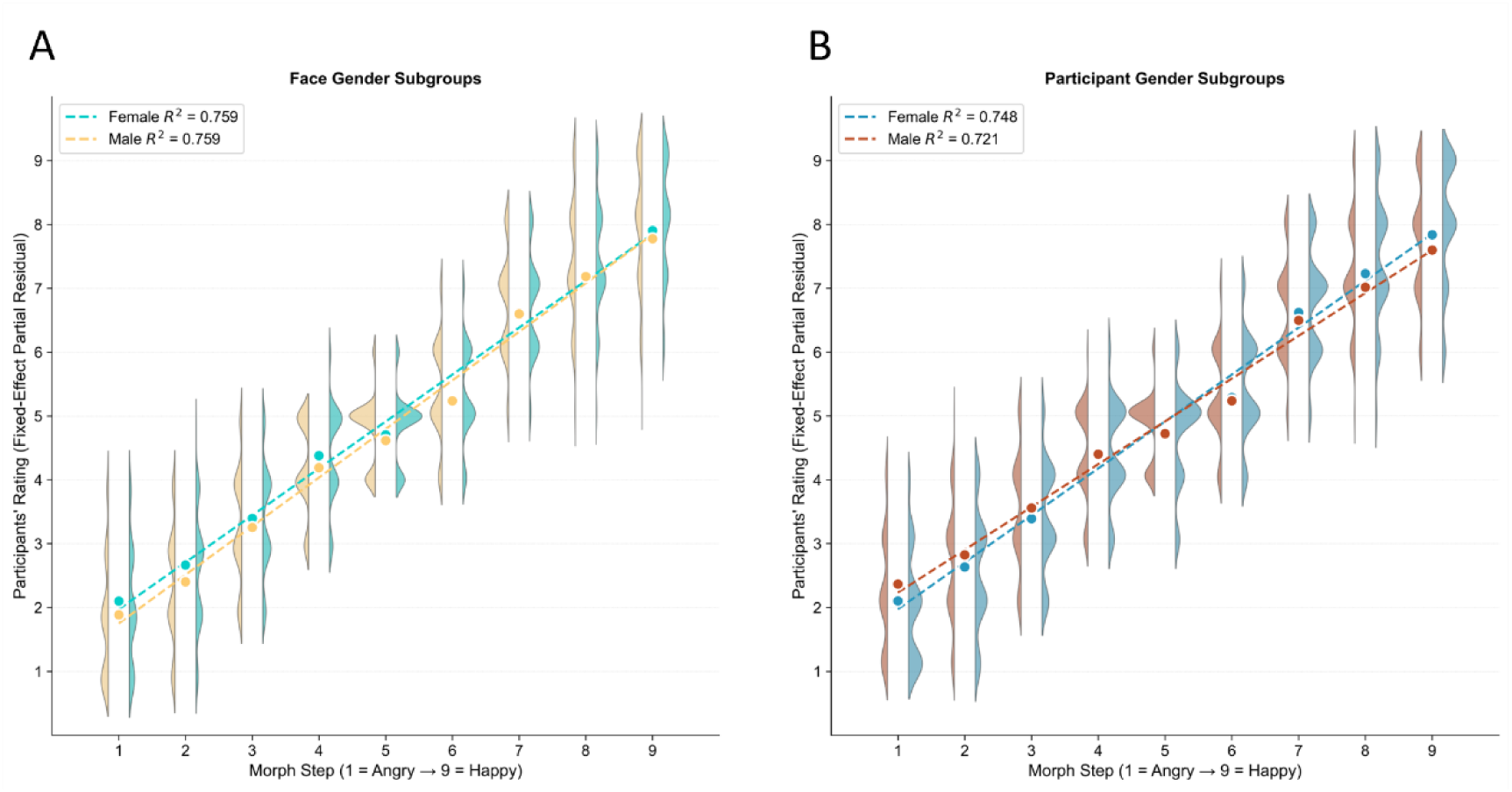
Gender-based subgroup analysis of the validity data. Note. A) Face-gender subgroups: female faces and male faces show comparable morph-rating slopes in model-derived partial residuals. B) Participant-gender subgroups: female participants show a steeper model-implied morph-rating slope than male participants. Violin plots display distributions of gender-specific partial residuals at each morph step (1: angry to 9: happy), with dashed lines showing mixed-effects-model-implied fixed-effect trendlines. Large markers indicate mean ratings per morph step. Extreme outliers (+/-2 SD from morph-level means) were excluded only for visualization clarity; all data were retained in the statistical models. These subgroup plots are descriptive and should be interpreted in light of the full mixed-effects model.

**Figure 6.**
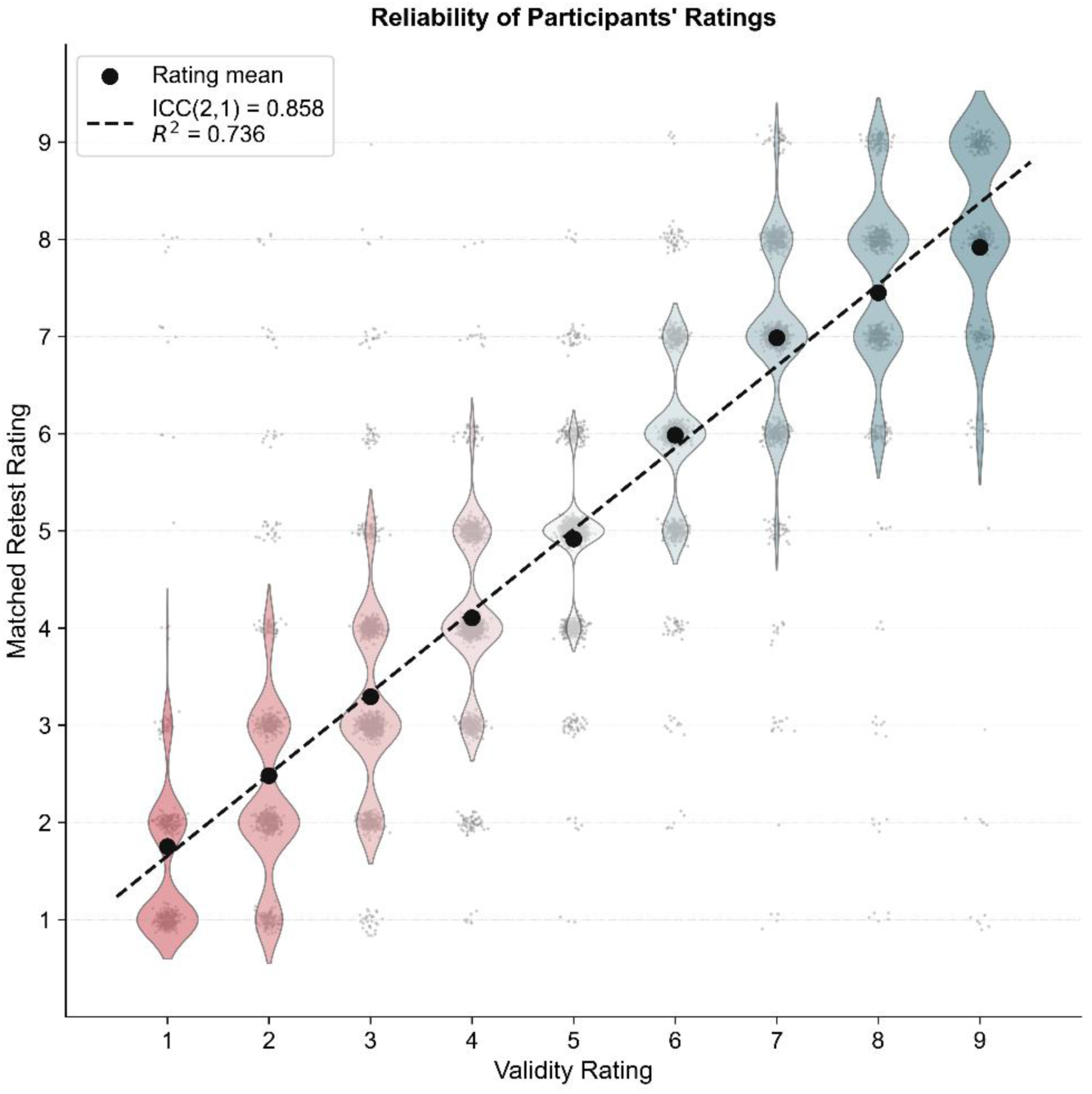
Reliability of participants’ ratings between the initial test and the retest session. Note. The x-axis denotes ratings from the first task, and the y-axis corresponds to matched ratings from the retest session. Violin plots depict the distribution and probability density of retest ratings at each initial rating level, with individual trials displayed by small grey dots. The means of the retest ratings are marked with black dots, and the dashed line shows the regression fit. The primary reliability estimate was ICC = 0.858, 95% CI [0.85, 0.86], indicating high absolute agreement across sessions. The regression R2 is reported only as a companion association-based summary.

For descriptive visualisation of subgroup patterns, we calculated partial residuals from the full model as described above. The model-implied fixed-effect lines for face-gender subgroups were:

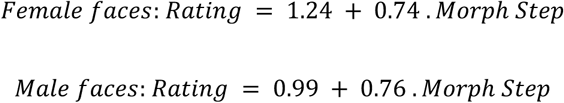

Thus, male faces showed a slightly steeper model-implied morph-rating slope, but the face-gender main effect was not significant once face identity was included as a random effect. The model-implied fixed-effect lines for participant-gender subgroups were:

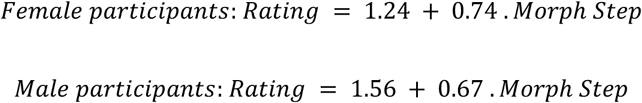

This pattern indicates that male participants tended to give higher ratings overall at the lower end of the continuum but showed a shallower morph-rating slope than female participants. These subgroup findings support the validity of the morph continuum across gender dimensions while also showing that gender-related patterns should be interpreted cautiously.

To evaluate whether the face-gender pattern reflects a broad, identity-consistent phenomenon or is driven by specific model faces, mean emotional ratings were calculated for each of the 22 *NimStim* identities by averaging across all nine morph steps and all participants (see Supplementary Figure 1a). Female identities produced a mean of 4.92 (SD = 0.16; range: 4.62-5.14), while male identities produced a mean of 4.79 (SD = 0.12; range: 4.59-4.92). Of the 11 female identities, 8 (73%) were rated above the male mean. A one-tailed Mann-Whitney U test confirmed a significant directional effect, U = 87.00, *p* = 0.0439, Cohen’s d = 0.875 (Supplementary Figure 1b). The one-tailed test was used because prior work motivated an a priori directional expectation that female faces would be rated as more emotionally expressive or happier than comparable male faces.

### Reliability

Test-retest reliability was assessed in the subgroup of 23 participants who completed the same task two weeks later. The primary reliability metric, ICC(2,1), indicated high absolute agreement between the initial and retest sessions (ICC = 0.858, 95% CI [0.85, 0.86]). As a companion association-based analysis, the regression of retest ratings on initial ratings produced a slope of 0.840, an intercept of 0.814, and R^2^ = 0.736. Together, these results suggest that ratings were stable across sessions within the retested subgroup.

## Discussion

In this paper, we introduced *STEMorph*, a newly developed stimulus set based on the *NimStim* dataset, featuring facial expressions transitioning from anger to happiness. We employed neutral-anchored morphing and neural-network-generated masking and evaluated the validity and reliability of this dataset.

Participants’ ratings tracked the intended morph continuum closely. In the mixed-effects model, morph step strongly predicted emotional ratings after accounting for repeated observations within participants and face identities. This supports the perceptual coherence of the anger-to-happiness continuum. However, as we did not directly compare *STEMorph* with alternative morphing methods, we cannot conclude that it produces more naturalistic stimuli than other morphing pipelines.

The gender-related findings provide useful normative information for future users of *STEMorph*, but they should be interpreted cautiously. Participant gender was associated with overall ratings and with the slope of the morph-rating relationship, and morph step interacted with face gender. These results align with previous studies suggesting that facial emotion recognition is influenced by the gender of the stimuli (23) and the perceiver (24, 36, 37). Prior research has consistently reported gender-based variations in emotion recognition (17) highlighting the importance of using stimuli that account for such differences. However, the main effect of face gender was not significant after accounting for face identity, indicating that identity-level variability contributes meaningfully to the observed pattern. Accordingly, these results should not be taken as strong evidence for a broad face-gender effect. Additionally, the high test-retest reliability observed in our study suggests minimal variability in interpretations across multiple exposures.

Our methodological choices also reflect advancements over traditional approaches. Specifically, utilizing a neutral expression as an intermediary anchor point minimized bias at the central position and facilitated smoother transitions between angry and happy expressions. This approach aligns with findings from earlier studies exploring emotional gradients (2) and builds on work demonstrating that neutral anchoring enhances naturalism in morphed stimuli (11). Perceptually, direct angry-to-happy morphs without a natural midpoint anchor tend to produce artefactual images at the transitional center, where opposing muscle-group activations — corrugator contraction associated with anger and zygomaticus major activation associated with happiness — must coexist within a single face. The result can appear uncanny rather than genuinely ambiguous, introducing a confound that is independent of the intended emotional content. This concern is supported by (11), who empirically demonstrated that the morphing strategy systematically influences emotion recognition bias, in part because perceptually incoherent midpoints distort the psychometric function. Conceptually, the neutral expression provides a semantically meaningful zero-point on the valence dimension, which is particularly valuable in bias research where the interest is the direction and magnitude of displacement relative to an emotional baseline.

Additionally, by generating distinct neural-network-based masks for each image, we mitigated issues like distortions in non-facial regions (e.g., hair or ears), which are common in datasets relying on simpler masking techniques (16). Moreover, while removing the background may reduce the real-world context of the stimuli, this step eliminates distractions, focusing solely on facial features for emotion recognition. These innovations ensure that *STEMorph* preserves anatomical fidelity while maintaining focus on facial features critical for emotion recognition.

Importantly, by making the *STEMorph* stimulus set publicly available along with all associated data and codes, we facilitate broader adoption and encourage further research in this critical domain of human interaction. This transparency aligns with calls within the literature for accessible resources to advance emotion recognition studies (9). Furthermore, *STEMorph’s* design principles could inform future efforts to create ecologically valid stimuli for diverse applications, including clinical diagnostics and machine learning algorithms for facial expression analysis.

Despite these strengths, this study has limitations that warrant consideration. While our strong linear relationships provide a promising foundation for future research, further examination across diverse samples is essential to assess the generalizability and external validity of our findings. For instance, expanding participant demographics beyond young adults of similar ethnic backgrounds could reveal how age or cultural differences influence emotion perception biases. Because the validation sample consisted of young adult medical students from Iranian institutions, the resulting normative data should be interpreted within this demographic and educational context, which may limit cross-cultural and cross-demographic generalisability.

Additionally, restricting participant evaluations to a scale from anger to happiness may simplify responses. Nonetheless, by focusing on the anger-happiness continuum, *STEMorph* provides a foundation for future studies exploring other emotional transitions such as sadness or fear, with more flexible response formats. Emotions such as sadness or fear may exhibit unique perceptual biases requiring tailored stimulus sets. Therefore, developing complementary transitions between various emotions could enrich our understanding of emotion recognition processes across different contexts. Furthermore, while the neutral-anchored morphing approach has clear perceptual advantages, it is worth acknowledging that anchoring to a neutral midpoint may reduce the degree of ambiguity at the central morph steps relative to a direct angry-to-happy continuum.

In conclusion, our results underscore the importance of methodological rigor in stimulus design within the field of emotion recognition. The incorporation of neutral-anchored morphing and neural-network-generated masks addresses limitations observed in earlier studies regarding the unnatural appearance of stimuli. By enhancing the naturalism and coherence of facial stimuli, our work significantly contributes to the ongoing efforts to improve methodologies in studying facial expressions. Furthermore, variations in emotional ratings across gender subgroups highlight the relevance of considering both participant and facial expressor characteristics in emotion recognition research. We believe that *STEMorph* can serve as a valuable resource for future studies exploring the dynamics of emotion perception across diverse populations. At the same time, the stimulus set should be interpreted within its intended scope: a controlled closed-mouth anger-to-happiness continuum validated in a specific sample. Future work should extend validation to additional emotions, more diverse populations, and direct comparisons with alternative morphing and masking approaches.

## Declarations

### Ethics approval

Approval was obtained from the ethics committee of Shahid Beheshti University of Medical Sciences (IR.SBMU.MSP.REC.1396.757). The procedures used in this study adhere to the tenets of the Declaration of Helsinki.

### Consent to participate

Informed written consent was obtained from all individual participants included in the study. Facial stimuli were reused from the *NimStim* Set of Facial Expressions (22), in which participants provided informed consent for the publication of their facial images, covering their reuse.

### Data and Code availability

The behavioural data and all code used for data acquisition and analysis in this study are openly available in the following GitHub repository: https://github.com/Mekatebi/STEMorph.

Researchers wishing to use the stimulus set should first request access to *NimStim* by following the instructions at https://danlab.psychology.columbia.edu/content/nimstim-set-facial-expressions. Once *NimStim* access has been granted, the *STEMorph* stimulus set is available from the corresponding author on reasonable request.

### Author Contributions

MEK and MHG performed research, MEK analyzed data, all designed research and wrote the paper.

---

**Supplementary Figure 1.**
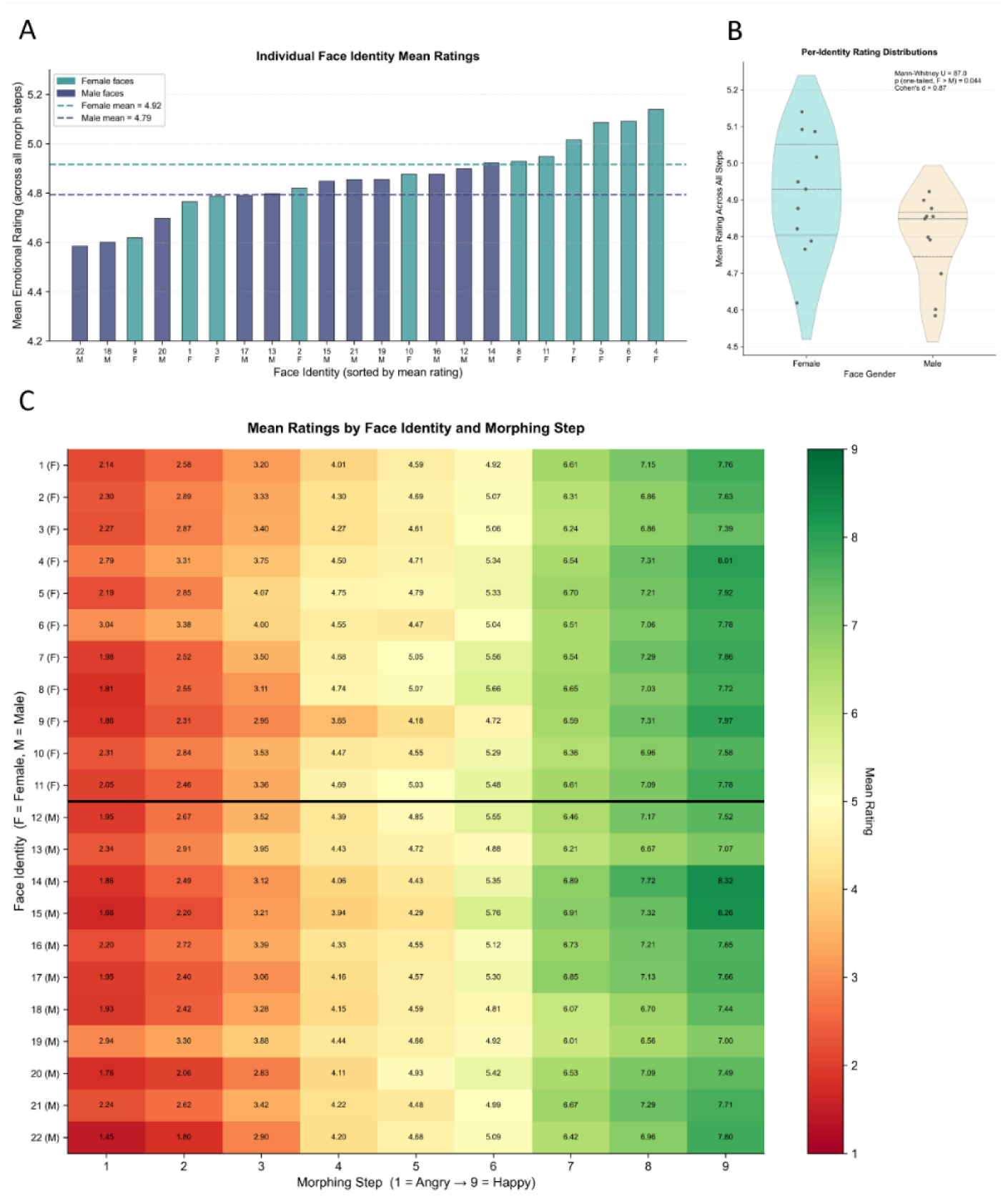
Face Identity Rating Profiles. Note. A) Mean emotional ratings averaged across all nine morph steps for each of the 22 face identities, sorted in ascending order. Female faces are shown in teal and male faces in navy. Dashed horizontal lines indicate the overall mean for female and male faces separately. Identity labels on the x-axis report the person number and gender abbreviation (F = Female, M = Male). B) Distribution of mean emotional ratings for female (teal) and male (navy) face identities. Violin shape represents the kernel density estimate. Individual face identity means are overlaid as jittered points. The red horizontal line within each violin marks the group median. Statistical comparisons: Mann–Whitney U test p-value (one-tailed, female > male), and Cohen’s d effect size. C) Mean emotional ratings for each of the 22 NimStim face identities (rows) across the nine morph steps (columns). Female faces (F, persons 1–11) are shown in the upper section and male faces (M, persons 12–22) in the lower section. Cell color reflects mean rating magnitude on a red–yellow–green scale; cell values are rounded to two decimal places.

